# Convective forces increase rostral delivery of intrathecal antisense oligonucleotides in the cynomolgus monkey nervous system

**DOI:** 10.1101/2020.03.26.008425

**Authors:** Jenna M. Sullivan, Curt Mazur, Daniel A. Wolf, Laura Horky, Nicolas Currier, Bethany Fitzsimmons, Jacob Hesterman, Rachel Pauplis, Scott Haller, Berit Powers, Leighla Tayefeh, Bea DeBrosse-Serra, Jack Hoppin, Holly Kordasiewicz, Eric E. Swayze, Ajay Verma

**Affiliations:** Ionis Pharmaceuticals, Inc., Carlsbad CA, USA; Biogen, Inc., Cambridge MA, USA; Invicro, LLC, Boston MA, USA; MPI Research, Mattawan MI, USA

**Author notes:** Co-first authors (Jenna M. Sullivan, Curt Mazur). Correspondence and requests for materials should be addressed to: Curt Mazur, Ionis Pharmaceuticals, Inc., 2855 Gazelle Court, Carlsbad, CA 92010.

**Keywords:** Intrathecal, lumbar puncture, convective forces, antisense oligonucleotide, PET imaging, SPECT imaging, CT imaging

## Abstract

**Background:** The intrathecal (IT) dosing route introduces drugs directly into the CSF to bypass the blood-brain barrier and gain direct access to the CNS. We evaluated the use of convective forces acting on the cerebrospinal fluid as a means for increasing rostral delivery of IT dosed radioactive tracer molecules and antisense oligonucleotides (ASO) in the monkey CNS. We also measured the cerebral spinal fluid (CSF) volume in a group of cynomolgus monkeys.

**Methods:** There are three studies presented, in each of which cynomolgus monkeys were injected into the IT space with radioactive tracer molecules and/or ASO by lumbar puncture in either a low or high volume. The first study used the radioactive tracer ^64^Cu-DOTA and PET imaging to evaluate the effect of the convective forces. The second study combined the injection of the radioactive tracer ^99m^Tc-DTPA and ASO, then used SPECT imaging and ex vivo tissue analysis of the effects of convective forces to bridge between the tracer and the ASO distributions. The third experiment evaluated the effects of different injection volumes on the distribution of an ASO. In the course of performing these studies we also measured the CSF volume in the subject monkeys by Magnetic Resonance Imaging.

**Results:** It was consistently found that larger bolus dose volumes produced greater rostral distribution along the neuraxis. Thoracic percussive treatment also increased rostral distribution of low volume injections. There was little added benefit on distribution by combining the thoracic percussive treatment with the high-volume injection. The CSF volume of the monkeys was found to be 11.9 ± 1.6 cm^3^.

**Conclusions:** These results indicate that increasing convective forces after IT injection increases distribution of molecules up the neuraxis. In particular, the use of high IT injection volumes will be useful to increase rostral CNS distribution of therapeutic ASOs for CNS diseases in the clinic.

## BACKGROUND

The development of novel CNS treatment modalities can be hampered by the lack of blood brain barrier (BBB) permeability. The intrathecal (IT) dosing route allows for direct delivery of therapeutic molecules to the central nervous system (CNS), bypassing the blood brain barrier. The most practical routine clinical route for IT drug delivery is via lumbar puncture, which introduces drug into the lumbar CSF cistern at the caudal end of the neuraxis. IT administration has been successfully used to deliver the antisense oligonucleotide (ASO) therapeutic Spinraza to patients with Spinal Muscular Atrophy (1). Additionally, several other BBB impermeable therapeutic molecules in development, including proteins, nucleic acids, viral gene therapy vectors, stem cells and exosomes are pursuing use of the IT dosing route to target diseases of the CNS. Since these approaches aim to treat various diseases in both pediatric and adult patient populations that can be associated with significant variability in inter-subject anatomy and CSF volumes, common principles are needed to optimize the IT dosing procedure for broad neuraxial delivery. Approaches that increase neuraxial drug exposure after lumbar IT delivery will be useful as ASO therapies, and other intrathecally delivered modalities, advance into indications that involve more rostral brain structures.

Broad CNS tissue drug delivery following IT dosing requires neuraxial movement within the subarachnoid CSF followed by entry into the CNS parenchyma (2). Drug movement within the subarachnoid space is largely facilitated by several CSF convective mechanisms including CSF turnover, cardiac motion, respiratory thoracic motion, and body movement (3-6). We recently demonstrated in rodent studies that convective force generated in the CSF by the IT dosing procedure itself can be leveraged to increase the spread of drug along the neuraxis (7). Here, we evaluate the convection enhancing effects of different dosing bolus volumes and externally applied percussive force on the neuraxial distribution and pharmacodynamic effect of ASOs delivered via IT lumbar puncture in cynomolgus monkeys with the goal of enhancing the cranial distribution and widespread CNS action of therapeutic ASOs.

To evaluate the effects of these convection inducing factors on IT drug delivery in a larger species, we conducted a series of experiments in non-human primates following IT delivery of imaging agents and ASOs. First, we quantified neuraxial distribution after IT drug delivery by PET imaging of the small molecule ^64^Copper-dodecane tetraacetic acid (^64^Cu-DOTA) (468 g/mole molecular weight), both with and without the use of convective forces. We then replicated these efforts with an ASO that was co-injected with ^99m^Tecnicium diethylenetriaminepentaacetic acid (^99m^Tc-DTPA) to allow for SPECT imaging to define the early distribution effects of the convective forces. Then the distribution, pharmacokinetics and pharmacodynamics of the ASO were evaluated and compared with the ^99m^Tc-DTPA imaging data. Finally, using a therapeutically relevant ASO targeting microtubule associated protein tau (MAPT), with the potential to treat Alzheimer’s disease and frontotemporal dementia (8), we confirmed that increasing convective forces can increase rostral CNS delivery of ASOs and imaging agents.

## METHODS

### Animals

Adult cynomolgus monkeys were used in the experiments described herein. Six were used for the ^64^Cu-DOTA experiment, 24 were used for the ^99m^Tc-DTPA/ASO experiment and 10 were used for the MAPT ASO experiment. The ^64^Cu-DOTA and ^99m^Tc-DTPA/ASO experiments were conducted at MPI Research (Mattawan, MI) and the MAPT ASO experiment was performed at Northern Biomedical Research (NBR, Norton Shores, MI). The 6 monkeys in the ^64^Cu-DOTA experiment were non-naïve, acquired from an existing colony at MPI Research (Mattawan, MI). The group was divided into 3 males and 3 females of 3.8 ± 0.7 kg body weight. The 24 animals in the ^99m^Tc-DTPA/ASO experiment were non-naïve and included 13 males and 11 females. Of this cohort, 20 animals received ^99m^Tc-DTPA/ASO and had a body weight of 4.1 ± 0.5 kg. The remaining four animals were kept naïve to ^99m^Tc-DTPA/ASO their CNS tissues were used as controls for the determination of metastasis associated lung adenocarcinoma (Malat1) RNA expression levels. For the Mapt experiment 10 animals were used, 5 female and 5 male, their body weights were between 3.0 and 4.5 kg. For all the animals, fluorescent lighting was provided via an automatic timer for 12 hours per day, tap water was supplied *ad libitum* via an automatic water system and they received an adequate supply of primate chow except during designated fasting periods.

### Anesthesia

Each animal was fasted for 4-12 hours prior to induction of anesthesia with ketamine (10-25 mg/kg. IM). Following anesthesia induction, animals were placed on isoflurane (1-2%) in oxygen carrier gas for the lumbar puncture, percussive wrap treatment when indicated, and imaging when indicated. Vital signs were monitored throughout the procedure. Lactated Ringer’s Solution was administered in a 5mL subcutaneous (SC) bolus both pre- and post-imaging. Each animal was also given intramuscular (IM) Metacam (0.2mg/kg, IM) and Ceftiofur (2.2mg/kg, IM).

### Magnetic Resonance Imaging (MRI)

Whole body MRI was acquired from each animal to measure CSF volume. All animals were imaged on a 1.5T Siemens Symphony MRI scanner (Siemens Medical Systems, Erlangen, Germany). High-resolution MRI of the spine and cranium were acquired in separate acquisitions within a single imaging session while each animal was under anesthesia. Images were acquired in axial orientation using a 3D T2 fast-spin echo (FSE) sequence with one excitation. Cranium data were acquired with a clinical CP extremity coil, voxel size of 0.7 × 0.7 × 2 mm, acquisition time of 8.05 min, TR of 12.5 ms, and TE of 6.25 ms. Body data were acquired with a clinical body coil, voxel size of 0.35 × 0.35 × 2 mm, acquisition time of 13.97 min, relaxation time of 11.82 ms, and excitation time of 5.91 ms.

### MRI data analysis

The cranium and body MRI data were stitched, including alignment, co-registration, and intensity blending in overlapping slices. Coarse, manually defined regions-of-interest (ROIs) were used to segment the brain and spine, including nearby surrounding tissue. Sub-regions with relative homogeneous CSF intensity within the brain and coarse spine ROI regions were selected. Sub-region-specific intensity thresholds were used to segment the final CSF ROI. In sub-regions with insufficient contrast, the CSF was segmented manually using MRI intensity as a guide.

### Lumbar Puncture Procedure

After each animal was anesthetized, an experienced veterinary surgeon injected the test articles to the intrathecal space by lumbar puncture. In the experiments performed at MPI, the animals were placed in left lateral recumbence and a 22-gauge Quinke spinal needle was introduced into the L4/L5 intrathecal space using aseptic technique. In the Mapt experiment performed at NBR, the animals were placed vertically in a seated position and the torso extended over a circular form. A 25-gauge Pencan Paed® pencil-point needle for pediatric use (B Braun) was inserted into the L4/L5 intervertebral space and used to inject the dosing solution. In all cases, the placement of the needle was verified by the presence of CSF at the needle hub pre- and post-injection. Once placement of the needle was verified, test article was injected over ∼2 minutes/mL volume, and then the needle was withdrawn.

### Percussive wrap and high frequency chest wall oscillation

In the ^64^Cu-DOTA and ^99m^Tc-DTPA/ASO experiments, a percussive wrap that delivers high-frequency chest wall oscillation (HFCWO) therapy (SmartVest^®^, Electromed, Inc., New Prague, MN) was applied to each animal after IT injection. The SmartVest^®^ system is an airway clearance system prescribed for patients with compromised airway clearance (such as in cystic fibrosis or spinal muscular atrophy). The wearable wrap consists of an inflatable bladder connected to an air pulse generating system. A size “S-Small” Single-Patient Use SmartVest Wrap^®^ with wrap height of 10.5 cm (4-1/4”) was used on all monkeys. Treatments were conducted at 5 Hz and 10 psi. Under percussive vest treatment conditions, animals were anaesthetized, test article was injected IT, then the wrap was applied to each animal’s torso. HFCWO treatment was applied for 30 minutes where indicated, otherwise the animal remained in the wrap for 30 minutes without activation of the air pulse mechanism. Following percussive wrap treatment, the wrap was removed, and each animal was placed in the PET or SPECT scanner for imaging.

### ^64^Cu-DOTA Radiochemistry

DOTA chelator was purchased from Macrocyclics (Lot M14010001-070713). ^64^Cu was supplied in 0.1M HCl by Washington University (St. Louis, MO). ^64^Cu-DOTA was prepared prior to each imaging session. Briefly, 0.1M, pH 6 citrate buffer, and ^64^Cu were added to a glass vial. Following the addition of radioactivity, chelator solution (1.0 mg/mL in 0.1M citrate buffer, pH 6) was added to the vial. The vial was vortexed for 1 min then placed in a water bath at 50°C for 30 min. Following chelation complex formation, two instant thin layer chromatography (ITLC) runs were conducted in parallel to confirm chelation efficiency.

For ITLC, a small aliquot of the ^64^Cu-chelator solution was diluted 1:100 in 200 µL. 2 µL were drawn up by pipette and dispensed onto the bottom of the ITLC paper strip and allowed to dry. Once dry, the ITLC strip was placed in a 15 mL conical tube containing 800 µL of ITLC developing buffer (0.1M citrate buffer, pH 6) and chelation efficiency was determined. Chelation efficiency was consistently >99%. Specific activity (SA) was 358.9 ± 166.5 MBq/mg (mean ± SD, n=8 runs).

### ^64^Cu-DOTA Study design

The tissue distribution of ^64^Cu-DOTA radioactivity was evaluated in 6 cynomolgus monkeys after IT bolus delivery of ∼18MBq ^64^Cu-DOTA under 4 conditions. Each condition consisted of an IT delivery of ^64^Cu-DOTA followed by dynamic, whole-body PET imaging. All animals were imaged under each condition in a crossover study design with 1-2 weeks between conditions. The tests on the different experimental days were low (0.36 mL) versus high (1.8 mL) injection volume, low injection volume versus low injection volume with 30-minute percussive wrap treatment, and high injection volume versus high injection volume with 30-minute percussive wrap treatment. There were no significant differences in molar mass of ^64^Cu-DOTA injected between conditions. Following the experiments, the animals were returned to the MPI colony.

### PET Image Acquisition

Whole-body continuous bed motion positron emission tomography (PET) and computed tomography (CT) data were acquired on a Focus220 microPET (Siemens Medical Systems, Knoxville, TN) and CereTom CT (NeuroLogica Corp, Danvers, MA), respectively. All animals were imaged in the head-first prone position. Radioactive fiducial markers were placed on the bed at three different positions for each scan to support co-registration of the PET time frames and as a standard of radioactivity. Dynamic PET data were acquired for either 0-120 minutes in scans without percussive treatment or 30-120 minutes in scans with percussive treatment PET data were acquired in 3D list-mode and re-binned into 2D sinograms. PET images were reconstructed by a 2D Ordered Subset Expectation Maximization (OSEM2D) algorithm with 14 subsets and 4 iterations. Corrections were made for detector normalization, decay, dead-time, random coincidences, and attenuation into images with 256×256×693 pixels and 0.95×0.95×0.80 mm voxels.

After each PET scan, a CT was acquired for anatomical registration. All animals were imaged on a specially design bed, which was transferred from the PET to the CT to allow consistent positioning of the animal between the two modalities. CT acquisition time was 6 minutes and based on an axial range of 450 mm, a tube peak voltage of 120 kVp, with 720 projections per rotation, and 4 seconds per projection.

### Estimation of tissue uptake of ^64^Cu-DOTA

ROIs were defined for the cranium, heart, liver, and kidneys by fitting ellipsoids of fixed volume to each organ based on CT. The cranium ROI contained both parenchyma and CSF within and surrounding the brain. The bladder ROI was defined through automated thresholding of the PET signal in the bladder or drawn by hand based on anatomy (in cases of low bladder signal). Spinal column ROIs were defined by applying a combination of manual and automated segmentation thresholds to the CT. The CSF ROI included both the spinal cord and CSF within the IT space. The CSF ROI was divided into cervical, thoracic, and lumbar sub-regions based on vertebral level.

Quantitative data from these ROIs were extracted and radioactivity concentration in units of percent injected dose per gram (%ID/g) at each time point for each ROI was plotted (assuming 1 cm^3^ is equivalent to 1 g of tissue). The area-under-the-curve (AUC) was calculated for each ROI. For AUC comparisons from conditions with and without percussive wrap treatment, AUCs for all regions were calculated from 30-120 min of data post-injection. Plots of radioactivity concentration (%ID/g) over the length of the spine were also generated for each ROI. All analyses were performed in VivoQuant™ 1.23patch1-3 and MATLAB R2014a (MathWorks®, Natick, MA, USA).

### ^99m^Tc-DTPA SPECT /ASO Study Design

Twenty cynomolgus monkeys were randomly assigned to 4 treatment groups of 5 each. All treatment groups received 16 mg of the ASO against *MALAT1* co-injected into the lumbar IT space with ∼37MBq of ^99m^Tc-DTPA. Two treatment groups received the ASO/tracer injection in a volume of 0.8 mL. These two groups were then divided into one with and one without 30 minutes of percussive wrap treatment. The other two treatment groups received the ASO/tracer injection in a volume of 2.4 mL. These two groups were also divided into one with and one without 30 minutes of percussive wrap treatment. An additional 4 animals naïve to any treatment were used as negative controls for the RNA expression and ASO concentration determinations.

### Antisense Oligonucleotide

The antisense oligonucleotide against both *MALAT1* and *MAPT* were developed by in vitro and in vivo screening techniques at Ionis Pharmaceuticals and were shown to be cross reactive to cynomolgus monkey (data not shown). These ASOs are both 20 base long chemically modified DNA molecules. The MALAT1 ASO has the sequence GCCAGGCTGGTTATGACTCA and the MAPT ASO the sequence ACACACCTTCATTTACTGTC. In both of the ASOs the 5 bases on both the 5’ and 3’ ends have 2’ methoxyethyl (MOE) sugar modifications and the backbones are mixtures of phosphothioate and phosphodiester linkages. Both ASOs were formulated in sterile artificial cerebral spinal fluid (aCSF). The Malat1 ASO was formulated at 20 and 6.7 mg/mL for doses of 16 mg in 0.8 and 2.4 mL, respectively. The MAPT ASO was formulated at 50 and 20 mg/mL for doses of 40 mg in 0.8 and 2.0 mL, respectively. Both ASOs were sterile filtered through a 0.22 um filter and placed in sterile septum vials which were crimped closed. The ASO vials were sent to the experimental sites for injection of the ASO into the animals.

### SPECT Image Acquisition

Immediately after IT ^99m^Tc-DTPA delivery, the animals were maintained under anesthesia and SPECT data were acquired using a SPECT/CT scanner (NanoSPECT/CT, Bioscan, Inc.) for 10 minutes (time 0 scan). Following this scan, each animal was placed in the percussive wrap for a 30 minute percussive wrap treatment (either activated or not) then imaged again with the SPECT scanner for 30 minutes (time 40 minute scan). The animals were allowed to recover from the anesthesia and then were re-anesthetized for a final 30 minute SPECT scan at 6 hours post IT injection. The animals were again allowed to recover from anesthesia. All SPECT scans were followed immediately by a CT scan.

The anatomical scan range was head to mid torso using a UHR parallel-hole collimator with 96 projections and an energy window of 126.5 – 154.6 keV. SPECT projection images were reconstructed in the software ReSPECT v 2.5 (SciVis, Germany) using a maximum likelihood estimation method (MLEM) with 6 iterations. Object background threshold was set to 5 and an attention coefficient of 0.12/cm was used. No smoothing was used in reconstruction and data were reconstructed into 1 mm isotropic voxels.

CT scans were performed from head to mid torso region on a CereTom CT (NeuroLogica Corp, Danvers, MA). Acquisition time was 604 seconds with tube peak voltage of 120 kVp, current set to 4 mA, 288 projections per rotation, and 6 seconds per projection.

### SPECT Image Processing and Image Generation

A single quantification factor (QF) was calculated and applied to each SPECT image. Three fiducial markers, consisting of vials containing a known amount of activity were included in each SPECT scan. Each fiducial marker was manually segmented from the unprocessed reconstructed SPECT image, and the given activity in that fiducial marker (decay-corrected to the scan time) was divided by the counts from the segmentation to obtain a QF for that fiducial. The QFs for the three fiducials were then averaged to obtain the QF for the image.

SPECT and CT images at each time point were co-registered. Maximum intensity projection (MIP) images of co-registered SPECT and CT were generated with SPECT data scaled from 0 to 1% ID/g. Image processing and MIP generation was performed in VivoQuant 2.0. Full quantitative analysis of the image data was not performed.

### ^99m^Tc-DTPA/MALAT1 ASO experiment - Euthanasia and Tissue Harvesting

One week (7 days) following the IT dosing procedure in the ^99m^Tc-DTPA/MALAT1 ASO experiment, the animals were euthanized by injection of euthanasia solution. Euthanasia was confirmed by the secondary method of exsanguination via femoral vessel incision. The brain, and spinal cord were harvested from the animals. The brains were placed in a monkey brain matrix (ASI Instruments, MBM-2500C) and sliced into 4 mm thick coronal slabs. Each of these slabs was divided into left and right hemispheres. The left hemisphere was immersion fixed in 10% phosphate buffered formaldehyde solution and the right hemisphere was frozen between Parafilm sheets on an aluminum plate cooled with dry ice and the tissue was buried under powdered dry ice. The spinal cord was divided into cervical, thoracic and lumbar sections from which small caudal pieces of the sections were immersion fixed in 10% phosphate buffered formaldehyde solution and the rostral pieces frozen buried in powdered dry ice. All frozen tissues were shipped to Ionis Pharmaceuticals for analysis of *MALAT1* RNA and tissue concentrations of the MALAT1 ASO. The fixed tissues were embedded in paraffin and histological sections were taken for immunohistochemistry (IHC) of the ASO and in situ staining of *MALAT1* RNA.

### MALAT1 RNA PCR Analysis

Tissue punches (∼12 mm^3^) were taken from fresh frozen brain and spinal cord slices and homogenized with sterile ceramic beads in guanidinium thiocyanate buffer containing 8% 2-mercaptoethanol using a bead homogenizer. Total RNA was prepared from tissue lysates using a RNeasy 96 kit (Qiagen). The prepared RNA was assayed for *MALAT1* and *cyclophilin A* levels using primer/TaqMan probe sets with an EXPRESS One-Step SuperScript quantitative reverse transcriptase polymerase chain reaction (qRT-PCR) kit (Invitrogen). qRT-PCR plates were run on StepOnePlus Real-Time PCR machines (Applied Biosystems) and data was initially analyzed using StepOne Software (Applied Biosystems). *MALAT1* RNA expression level was normalized to *cyclophilin A* mRNA expression to correct for the amount of RNA in the reaction. Malat1 RNA expression level was further normalized as percent naïve control.

Malat1 primers/probe sequences:

Forward 5’-AAAGCAAGGTCTCCCCACAA-3’

Reverse 5’-GTGAAGGGTCTGTGCTAGATC-3’,

Probe 5’-/56-FAM/CAACTTCTC/ZEN/TGCCACATCGCCACCT/3IABkFQ-3’;

Cyclophilin A primers/probe sequences:

Forward 5’-CGACGGCGAGCCTTTG-3’

Reverse 5’-TCTGCTGTCTTTGGAACCTTGTC-3’,

Probe 5’-/56-FAM/CGCGTCTCCTTCGAGCTGTTTGC/36-TAMSp/-3’

### Analysis of Tissue Levels of ION-626112

Frozen tissue samples from frontal cortex, lumbar, thoracic and cervical spinal cord were obtained from the frozen monkey tissues. Where possible the samples for PCR analysis and tissue concentrations of the Malat1 ASO were adjacent samples. The samples were minced, weighed into individual wells in a 96-well plate, and had 500µL homogenization buffer added to those wells. Control tissue homogenate for curves was made by weighing control monkey brain and adding homogenization buffer at a 9 to 1 ratio. Five hundred microliter aliquots were pipetted into a 96-well plate and appropriate amounts of calibration standards were spiked in the corresponding wells. Wells containing samples and calibration standards then had internal standard (IS) and approximately 0.25cm^3^ granite beads added. The plates were then extracted via a liquid-liquid extraction with ammonium hydroxide and phenol:chloroform:isoamyl alcohol (25:24:1). The aqueous layer was then further processed via solid phase extraction on a Strata X plate (Phenomonex Inc., CA). Eluates had a final pass through a protein precipitation plate before they were dried under nitrogen at 50°C. Dried samples were reconstituted in 140µL water containing 100µM EDTA. An Agilent liquid chromatograph with mass spectrometry detection (LC-MS/MS) instrument consisting of a 1290 binary pump, a column oven, an auto sampler, and a 6460 triple quadrupole mass spectrometer was used for analysis (Agilent, Wilmington, DE, USA).

The tissue extracts were injected onto a Kinetex analytical column (Phenomenex, 100 × 2.1 mm, 2.6 µm particle size; 100 Å) which was equilibrated with 15% methanol in 5mM triethyalamine (TEA) and 400mM hexo-fluoro-isopropanol (HFIP) and then maintained at 55°C and with a flow rate of 0.3 mL/min throughout the analysis. A gradient from 15 to 50% methanol over 7 minutes, increased to 80% over 0.5 minutes, and maintained at 80% for 0.5 minutes was used to separate the Malat1 ASO and internal standard (IS) from background peaks. A re-equilibration time of 2 minutes at the starting conditions was used between samples.

All mass measurements were made on-line with the scan time set from 2 to 8 minutes. During that time window, the mass spectrometer was set to scan for MRM transitions for the full length ASO and IS (−8m/z, 893.1->94.8; and -4m/z, 1160.7->94.8, respectively). Mass spectra were obtained using a spray voltage of -1500 V, a nebulizer gas flow of 25 psig, a sheath gas flow rate of 12 L/min at 350°C, a drying gas flow rate of 5 L/min at 250 °C, and a capillary voltage of –3750 V. Chromatograms were analyzed using Agilent Mass Hunter software.

Tissue calibration curves were constructed using peak area ratios of the ASO to the IS and applying a weighted (1/x) Quadratic regression. All tissues had a calibration range for the ASO from 0.036 - 178.82 µg/g in 50 mg monkey brain homogenate. A minimum signal-to-noise ratio of 5:1 was used to distinguish ASO peaks from background. Acceptance criteria for the calibration curve were set to 85-115% of nominal values. All samples were stored at -70°C ± 10°C, upon receipt.

### In situ Staining of MALAT1 RNA and IHC for MALAT1 ASO

Based on the PCR results, it was determined that animal numbers 1011 and 1006 were typical of the *MALAT1* knock down level of the successfully dosed animals in the high volume no wrap and low volume no wrap groups, respectively and were chosen for in situ staining for *MALAT1* RNA. Naïve animal 2601 was chosen as a normal control for the *MALAT1* in situ staining as comparator. The paraffin embedded tissues were sectioned at 4 um thickness and in the case of the tissue block being larger than the 1” x 3” slides they were sliced down the center with a razor blade and the two halves were collected onto separate slides. Adjacent sections were taken from all tissue blocks from animals 1011, 1006 and 2601 for the *MALAT1* in situ stain and immunohistochemistry for the ASO.

The in-situ stain for *MALAT1* RNA was performed on a Leica Bond RX research staining robot with a RNAScope custom reagent kit from Advanced Cell Diagnostics (ACD). A 2.5 LS reagent kit – Brown (ACD, #322100), a RM-Malat1 probe 01-04 (ACD, #460238), DapB negative control and PPIB positive control were employed for the stain. Initially, the slides were dried overnight and then again for 30 minutes in a 60°C oven. The slides were then loaded into the Bond RX machine and the following RNAScope 2.5 LS reagents were used in the automatic staining procedure: hydrogen peroxide, protease III, AMP1-AMP4, AMP5 and AMP 6 Brown, Rinse, bluing reagent, *MALAT1* target probe, Mock probe wash and Leica polymer refine detection kit. The automated run was completed then the slides were rinsed in distilled water, dehydrated through graded alcohol to a clearing agent, and cover slipped with Permount (Leica Micromount, #3801731). The slides were allowed to dry and then were scanned into a Leica Aperio scanner at 20X resolution.

For the immunohistochemical staining for the ASO, sections were cut from the paraffin embedded blocks at 4um thickness onto positive charged slides and dried overnight at 50°C. The tissues were deparaffinized to distilled water and the antigen retrieval performed with Proteinase K (Dako #S302030-2) treatment for 2 minutes. Endogenous peroxidase was quenched by incubation in Dako Dual Endogenous Enzyme Blocking Reagent (#S2003) for 10 minutes followed with a 30 minute incubation in Cyto-Q Background Buster (Innovex, #NB306-125ml) to block nonspecific protein binding. The slides were incubated with the custom made anti-ASO primary antibody, 6651 (Pan ASO, Ionis Pharmaceuticals) diluted 1:40,000 for 1 hour at room temperature. The slides were incubated for 30 minutes in a 1:200 dilution of the goat anti-rabbit secondary antibody conjugated to horse radish peroxidase (Jackson ImmunoResearch #111-036-003). The brown color was produced by reaction with 3,3’-diaminobenzidine chromogen (DAB) (Dako, #K3468) for 5 minutes and the tissues were counterstained with hematoxylin, dehydrated, cleared and cover slipped with Leica Micromount. After drying, the slides were scanned into the Hamamatsu Nanozoomer scanner at 20X resolution.

The scanned slides were visualized on a computer and images from the different CNS anatomical regions were acquired by screen capture.

### MAPT ASO experimental design

A group of 7 cynomolgus monkeys was randomly assigned to two groups of 5 for ASO treatment and one group of 2 for vehicle treatment. Each of the two groups of 5 were treated by intrathecal lumbar puncture with 40 mg of the MAPT ASO in either a low (0.8 mL) or high (2 mL) dosing volume every two weeks for 4 doses (on days 1, 14, 28 and 56). The vehicle treated animals were treated with 2 mL of aCSF on the same dosing schedule. All doses were delivered by hand at an approximate rate of 2 mL/minute. The animals were euthanized 7 days following the final dose (on day 63), brain and spinal cord were harvested and frozen for analysis of *MAPT* mRNA expression and ASO tissue concentration. Samples of the lumbar spinal cord, thoracic spinal cord, and frontal cortex, were dissected from the frozen tissue samples for *MAPT* mRNA expression analysis by qRT-PCR and adjacent samples were collected for determination of MAPT ASO concentrations.

## Statistics

In the ^64^Cu-DOTA experiment area under the curve calculations for radioactivity versus time were performed using the trapezoidal rule using GraphPad Prism (version 8.0.2). Comparisons between treatment modalities (high versus low volume, low volume versus low volume with wrap treatment, high volume versus high volume with wrap treatment) were done with two-tailed Welch’s T-tests using GraphPad Prism (version 8.0.2). In the ^99m^Tc-DTPA/ASO experiment differences between treatment groups in each tissue for MALAT1 ASO concentration and *MALAT1* RNA expression were evaluated with Two-way ANOVAs with Tukey’s multiple comparisons test using GraphPad Prism (version 8.0.2). In the MAPT ASO experiment differences between low and high dose volume for MAPT ASO concentrations and MAPT RNA expression were done with two-tailed Welch’s T-tests using GraphPad Prism (version 8.0.2). When errors are reported they are standard deviations. For all comparisons, P < 0.05 was considered statistically significant. The coefficient of variation was calculated by the formula: Standard deviation/mean*100%.

## RESULTS

### MRI determination of CSF volume in Cynomolgus monkeys

To support the selection of IT dose volumes, total CSF volumes were measured in cynomolgus monkeys. The CSF volume in 20 animals (10 males and 10 females, 3.6 ± 0.4 kg body weight, no significant differences between male and female body weights), was 11.6 ± 1.5 cm^3^ for the entire neuraxis including the ventricles (Table 1). Total CSF volume was 12.1 ± 1.6 cm^3^ in males and 11.0 ± 1.1 cm^3^ in females (Table 1). There were no statistically significant differences between males and females in CSF volume (Table 1).

**Table 1.** Whole body CSF volume determined in cynomolgus monkeys using MRI image analysis.

| Animal # | Sex | Body Weight (kg) | Total CSF volume (cm<sup>3</sup>) |
| --- | --- | --- | --- |
| A1001 | male | 3.7 | 14.4 |
| A1002 | female | 3.1 | 8.8 |
| A1003 | male | 3.3 | 11.0 |
| A1004 | female | 3.3 | 10.0 |
| A1005 | male | 3.7 | 10.3 |
| A1006 | female | 4.1 | 11.8 |
| A1007 | female | 3.8 | 11.9 |
| A1008 | male | 4.2 | 11.7 |
| A1009 | female | 3.1 | 12.6 |
| A1010 | male | 3.4 | 12.4 |
| A1011 | female | 2.8 | 10.4 |
| A1012 | male | 3.7 | 13.2 |
| A1013 | female | 3.2 | 10.6 |
| A1014 | male | 3.1 | 10.4 |
| A1015 | female | 4.0 | 12.3 |
| A1016 | male | 4.2 | 11.1 |
| A1017 | female | 4.1 | 10.9 |
| A1018 | male | 3.2 | 11.5 |
| A1019 | male | 4.3 | 14.9 |
| A1020 | female | 3.4 | 11.0 |
| Analysis of Total CSF volume (cm<sup>3</sup>) |  |  |  |
| Parameter | All Animals | Female | Male |
| Mean | 11.6 | 11.0 | 12.1 |
| SD | 1.5 | 1.1 | 1.6 |
| CV% | 12.7% | 10.4% | 13.3% |
| Median | 11.3 | 10.9 | 11.6 |
| Max | 14.9 | 12.6 | 14.9 |
| Min | 8.8 | 8.8 | 10.3 |
| Range | 8.8-14.9 | 8.8-12.6 | 10.3-14.9 |
| Female vs. Male T-test |  | 0.1043 |  |

To aid in translation of this work, we chose the experimental dosing volumes in NHP to be a similar percentage of total CSF volume as dose volumes used previously in human patients. High intrathecal dose volumes used in humans have corresponded to approximately 14% of total adult human CSF volume and volumes of up to 33% of the total CSF volume have previously been given safely in humans (9-11). We therefore evaluated the effect of IT dose volumes ranging from ∼3%- 20% of total monkey CSF volume determined by MRI with the amount of injected molecule kept constant within an experiment. Specifically, IT bolus injection volumes representing ∼3% (0.36 mL) and 15% (1.8 mL) of the total CSF volume were used in the ^64^Cu-DOTA experiment. For the ^99m^Tc-DTPA/Malat1 ASO experiment, IT bolus injection volumes representing ∼7% (0.8 mL) and 20% (2.4 mL) of total CSF volume were injected, and for the MAPT ASO experiment IT bolus volumes representing ∼7% (0.8 mL) or high injection volume representing ∼17% (2 mL) of total CSF volume were injected

### Exclusion of Animals from data analysis

In some cases, in the ^64^Cu-DOTA and the ^99m^Tc-DTPA/ASO experiments it was determined that there were IT doses missed due to technical failures. These animals were removed from further analysis. The following are the criteria used for the exclusion of these animals.

In the ^64^Cu-DOTA experiment, the surgeon assessed all IT injections to be successful, except for the injection for one subject under the low volume plus wrap condition. Despite the apparent success of the injections, the image data suggested that at least some injections were multi-compartmental (e.g. IT and epidural space) or only partially in the IT space (Figure 1A). A criterion to exclude poor or partial IT-injections was defined based on image features: images in which there was an obvious subdermal injection, a non-continuous signal in the IT space or a strong kidney signal at the first imaging time point after injection (reflecting early leakage to the periphery) were excluded from subsequent analyses. Six scans were excluded from analysis based on these criteria: two from the low volume group, one from the high-volume group, two from the low volume plus wrap group, and one from the high volume plus percussive wrap group.

**Figure 1:**
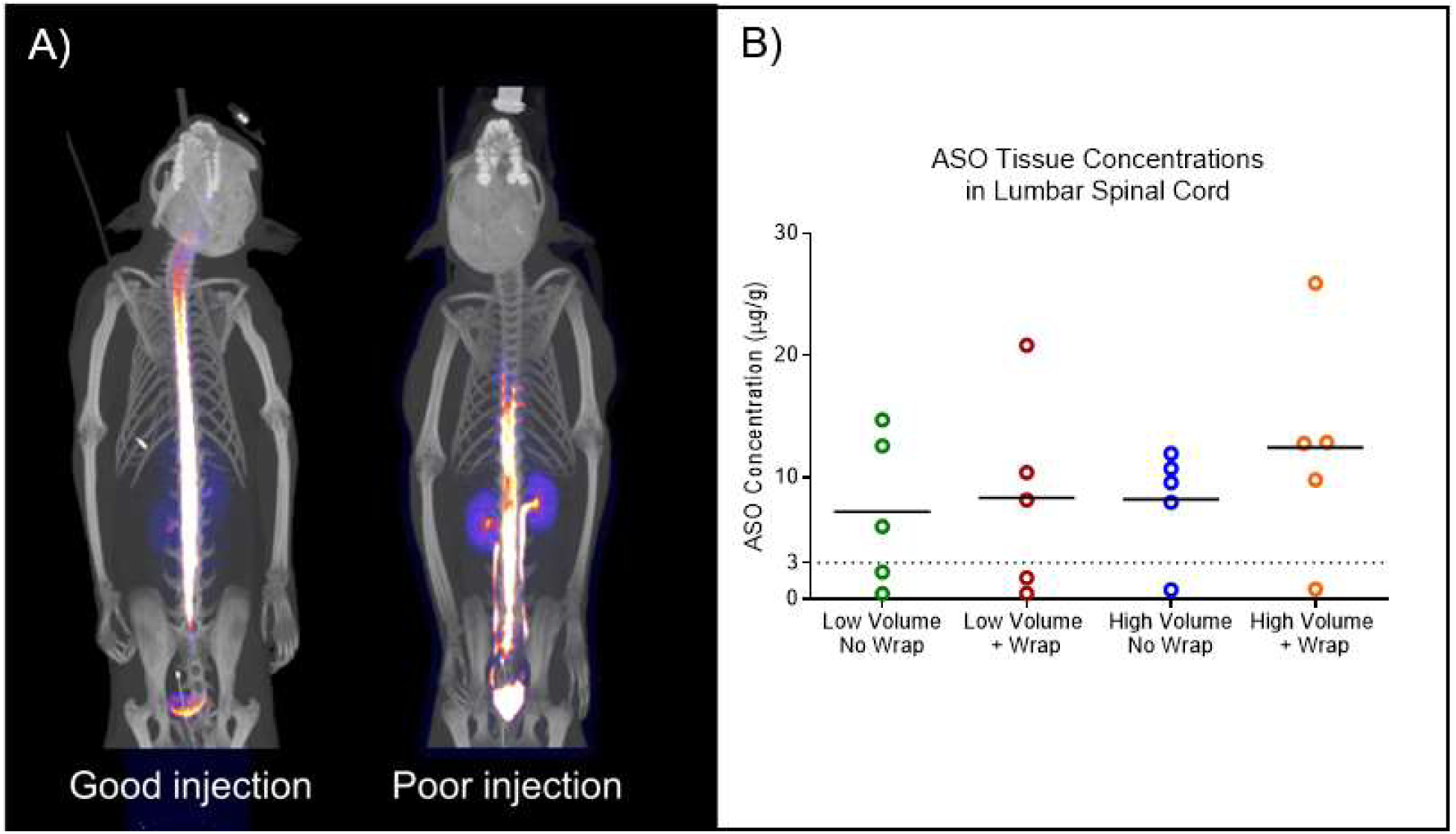
A) Imaging rejection criteria for poor technical injections in the ^64^Cu-DOTA experiment. The PET/CT scan on the left is indicative of a successful IT injection where the radioisotope is mostly contained in the neuraxial compartments. The PET/CT scan on the right illustrates an unsuccessful IT injection where a high radioisotope signal can be seen in the kidneys, ureters and bladder. B) Graph of the ASO tissue concentrations in µg/g tissue in the lumbar spinal cords of all the animals. Animals with lumbar spinal cord tissue concentrations below 3 µg/g tissue were considered probable missed injections due to technical reasons. There were two animals in both low volume (0.8 mL) groups and one animal in each high dose volume (2.4 mL) group that had lumbar spinal cord concentrations below 3 µg/g.

Because of the limitations of the field of view for the SPECT system used in the ^99m^Tc-DTPA/ASO study, the lumbar spinal area could not be imaged and so the success of the injections could not be assessed as was done with the ^64^Cu-DOTA experiment. Instead, we used the tissue concentrations of the co-injected ASO in the lumbar spinal tissue to determine technical success of the IT injections. From the graph of the lumbar spinal cord ASO concentrations in Figure 1B, it is quite apparent that two animals from group injected with low volume without percussive wrap, two animals from low volume plus percussive wrap group, one animal from high volume without percussive wrap group and one animal from the high volume with percussive wrap group had very low tissue concentrations when compared to the other animals in these groups (<3 ug/g). These six animals were termed technical missed dose animals and removed from further analysis.

### Increasing dose volume or adding external percussion increases rostral ^64^Cu-DOTA distribution along the neuraxis

To determine if drug volume or percussion could alter distribution of an IT delivered drug, ^64^Cu-DOTA was injected IT into cynomolgus monkey. The compound was delivered under four different paradigms, low volume (0.4 mL, 3% of CSF volume) without percussion, high volume (1.8 mL, 15% of CSF volume) without percussion, low volume with percussion, and high volume with percussion. These four paradigms were evaluated in three different experiments using the same 6 animals: the first evaluated low versus high injection volume, the second evaluated low volume versus low volume with percussion and the third evaluated high volume versus high volume with percussion.

Basic in-life tolerability assessments were included to confirm the tolerability of the manipulations, and that they did not alter vital signs that may contribute to CSF dynamics. The test article, ^64^Cu-DOTA, was well-tolerated. Percussion treatments, delivered using a percussive wrap and HFCWO, were also well-tolerated. Vital signs were within normal ranges for cynomolgus monkeys: heart rate was 132 ± 23 BPM, respiratory rate was 31 ± 8 respirations/min, and body temperature was 96.9 ± 1.9 °F.

The AUC of the ^64^Cu-DOTA was observed in the lumbar spine near the injection site with the concentration decreasing along a caudal to rostral gradient. Comparing the low and high-volume IT injection conditions, there were no significant differences between the ^64^Cu-DOTA AUC values in the lumbar spinal cord, the site nearest to the injection, of the two volumes (Figure 2A). The AUC values in the cranium and the thoracic spine were significantly higher in the high-volume condition than the low volume condition. There was also a nonsignificant trend towards higher cervical spinal concentrations with the higher volume compared to the low volume injection (P = 0.0624).

**Figure 2:**
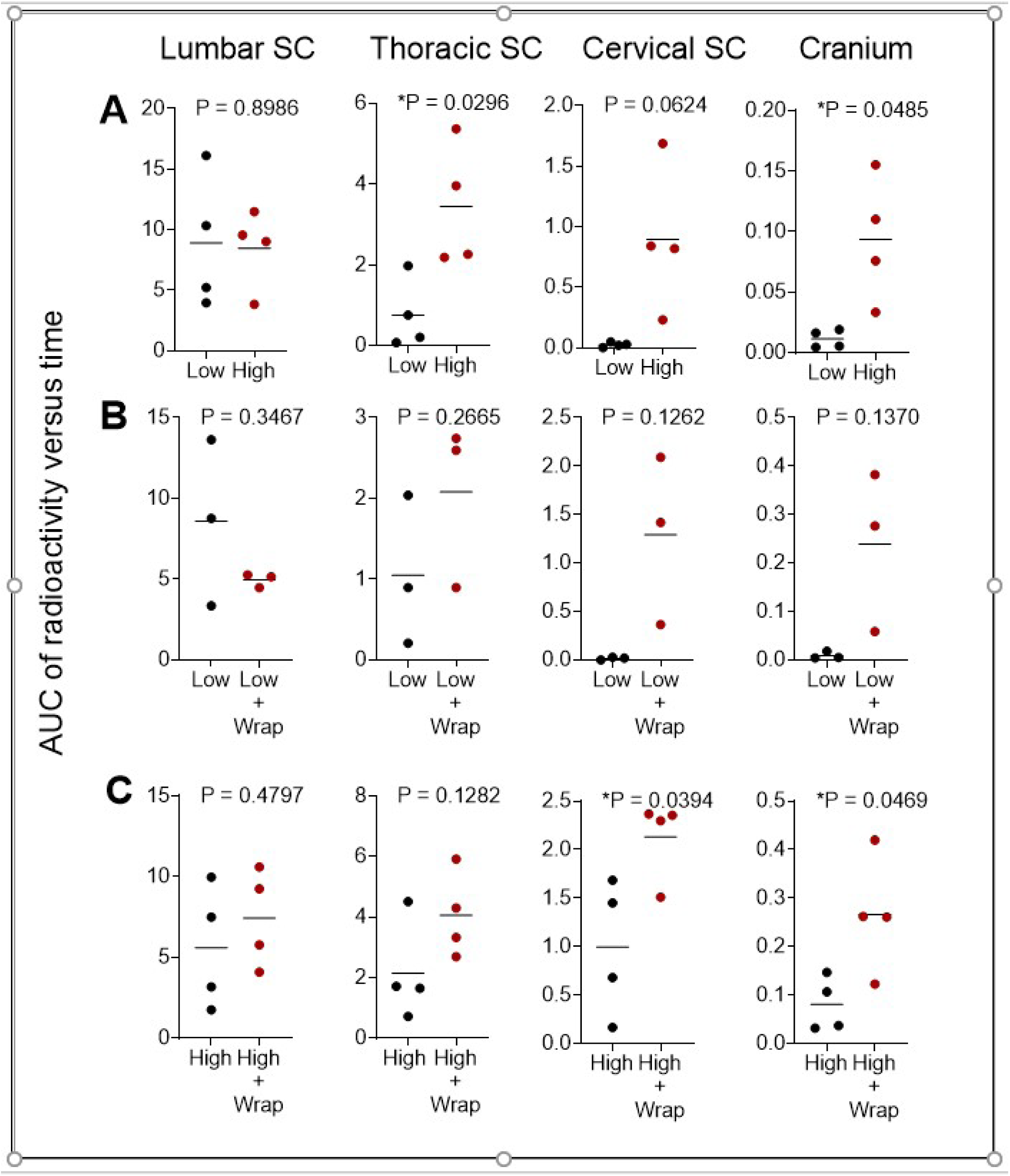
Area under the curve analysis (AUC) of the radioactivity concentration versus time curves from the lumbar, thoracic, and cervical spinal regions and the cranial region of the PET scans. AUC values are in units of %ID/g * hr. A) Comparisons of low (0.4 mL) versus high (1.8 mL) injection volume in the different CNS tissues. B) Comparisons between low injection volume versus low injection volume with percussive wrap treatment in the different CNS tissues. C) Comparisons between high injection volume and high injection volume with percussive wrap treatment in the different CNS tissues. Data displayed are individual values with a horizontal bar depicting the mean for each group. The P values resulting from the two-sided Welch’s T-test are written above each graph and a star (*) designates a P value less than 0.05.

To determine if percussive wrap treatment could further influence distribution, we compared low and high volumes of ^64^Cu-DOTA with percussive wrap treatment (Figure 2B and 2C, respectively). With low volume IT injections, percussive wrap treatment resulted in a trend towards greater rostral distribution of the ^64^Cu-DOTA, with a clear trend for higher AUC values in the more rostral cervical spine and cranial regions with wrap treatment (Figure 2B). With high-volume IT injections, percussive wrap treatment also resulted in greater rostral distribution of the ^64^Cu-DOTA, significantly increasing rostral distribution of the high volume ^64^Cu-DOTA to the cervical spine and cranial regions (Figure 2C).

### Increasing dose volume or adding external percussion increases rostral ^99 m^Tc-DTPA and ASO distribution along the neuraxis

To determine if we could replicate the results from the ^64^Cu-DOTA experiment with a therapeutically relevant modality, ASOs, and extend the findings to further understand the kinetics of distribution, we co-injected ASO with ^99m^Tc-DTPA. There were again four different paradigms, low volume (0.8 mL, 7% of CSF volume) without percussive wrap, high volume (2.4 mL, 20% of CSF volume) without percussive wrap, low volume with percussive wrap, and high volume with percussive wrap. These different paradigms were tested in the same experiment using different groups of monkeys for each treatment.

After IT co-injecting ^99m^Tc-DTPA with MALAT1 ASO the animals were imaged using SPECT at 0 min (prior to percussive wrap treatment), 40 min (after percussive wrap treatment) and 6 hours post-injection. The images were limited to the cranial, cervical, and thoracic spinal cord due to limitations of the SPECT system’s field of view. In the low injection volume with no percussive wrap group (Figure 3A) the radioisotope progressed up the spinal column. Addition of percussive wrap treatment to the low dose volume (figure 3B) increased the rostral distribution of the radioisotope signal into the cervical spine and brainstem area. Increasing the dose volume from low to high with no percussive wrap treatment (Figure 3C) resulted in greater rostral distribution in the base of the brain and into the transverse fissure between cerebellum and cerebrum and the lateral fissure between the temporal and frontal lobes. Addition of percussive wrap treatment to the high dose volume did not appreciably affect the distribution of the radioisotope over high volume alone (Figure 3D).

**Figure 3:**
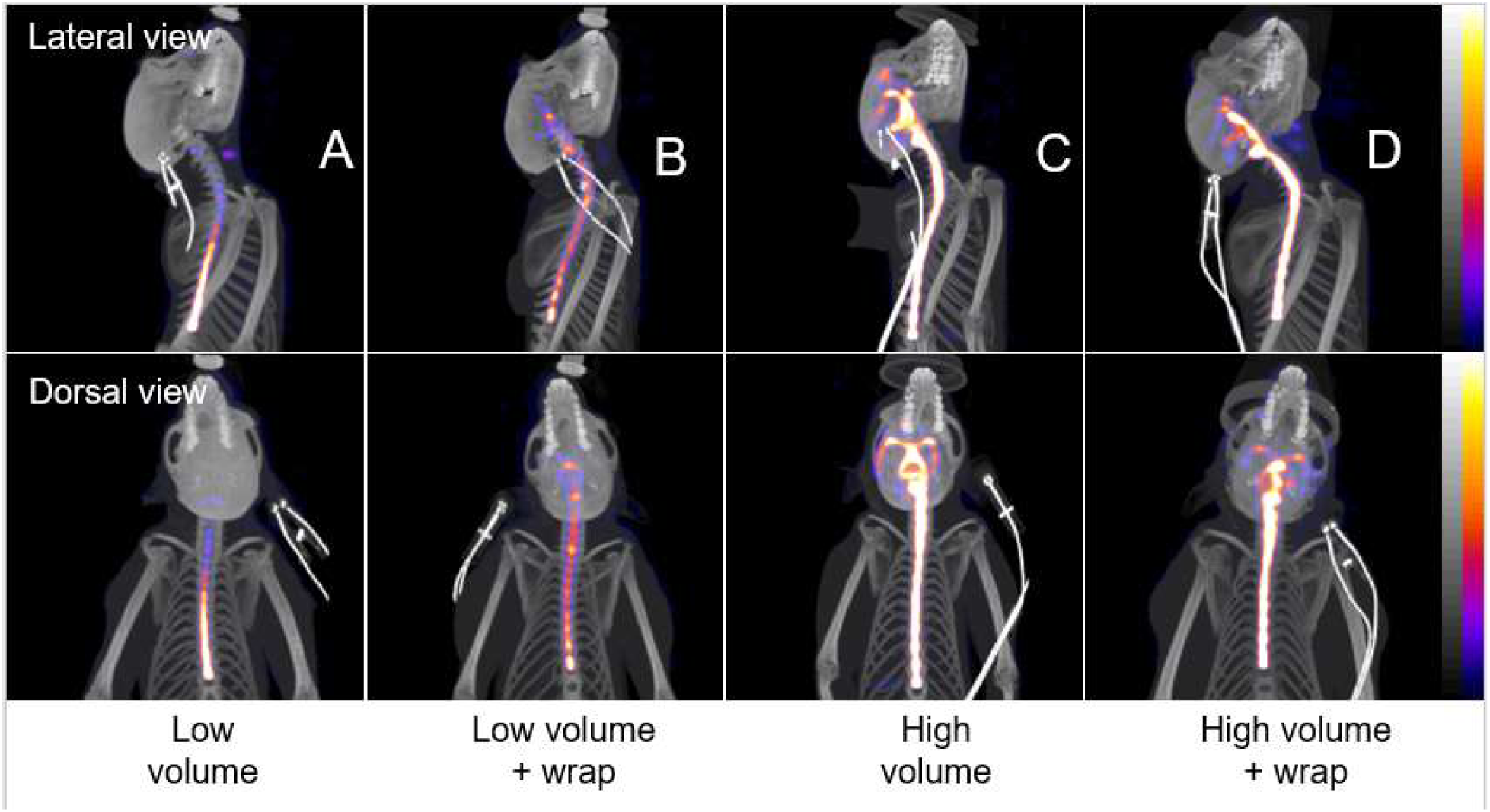
Co-registered maximum intensity projection SPECT and CT images at 40-minutes post-injection from representative animals from the low (0.8 mL) volume (A), low volume plus percussive wrap (B), high (2.4 mL) volume (C), or high volume plus percussive wrap (D) treatment groups. The ^99m^Tc-DTPA tracer incorporated into the dosing solution appears as bright areas within the spinal and cranial areas. SPECT images are scaled from 0 to 1% injected dose/g.

To determine the distribution of ASO into tissues, samples from the lumbar, thoracic, cervical spinal cord, and frontal cortex were harvested 7 days after injection and analyzed for concentrations of ASO. Tissue concentrations across the CNS are lower in the low volume group (Figure 4A) than those in the tissues of the animals dosed with the high dose volume (Figure 4C). Consistent with the SPECT imaging (Figure 3), the rostral neuraxial distribution appeared to be increased by increasing the dose volume, where the percussive wrap treatment was beneficial in the low dose volume group (Figure 4B versus 4A), but in this experiment, not in the high dose volume groups (Figure 4D versus 4C). When the ASO concentrations in the different CNS tissues were analyzed across treatments, no significant differences were detected, however there was a trend towards higher tissue concentrations in the Frontal cortex when low volume versus high volume were compared (Table 2).

**Table 2.** Tukey’s multiple comparisons test for ASO concentration results

| Within each row, compare columns<br>(simple effects within rows) | Adjusted P Value |  |  |  |
| --- | --- | --- | --- | --- |
|  | Lumbar<br>Spinal<br>Cord | Thoracic<br>Spinal<br>Cord | Cervical<br>Spinal<br>Cord | Frontal<br>Cortex |
| Low Volume no Wrap vs.<br>High Volume No Wrap | 0.9818 | 0.8036 | 0.7176 | 0.1114 |
| Low Volume no Wrap vs.<br>Low Volume + Wrap | 0.8994 | 0.4168 | 0.9976 | 0.7696 |
| High Volume No Wrap vs.<br>High Volume + Wrap | 0.1791 | 0.9998 | 0.9807 | 0.4357 |

**Figure 4:**
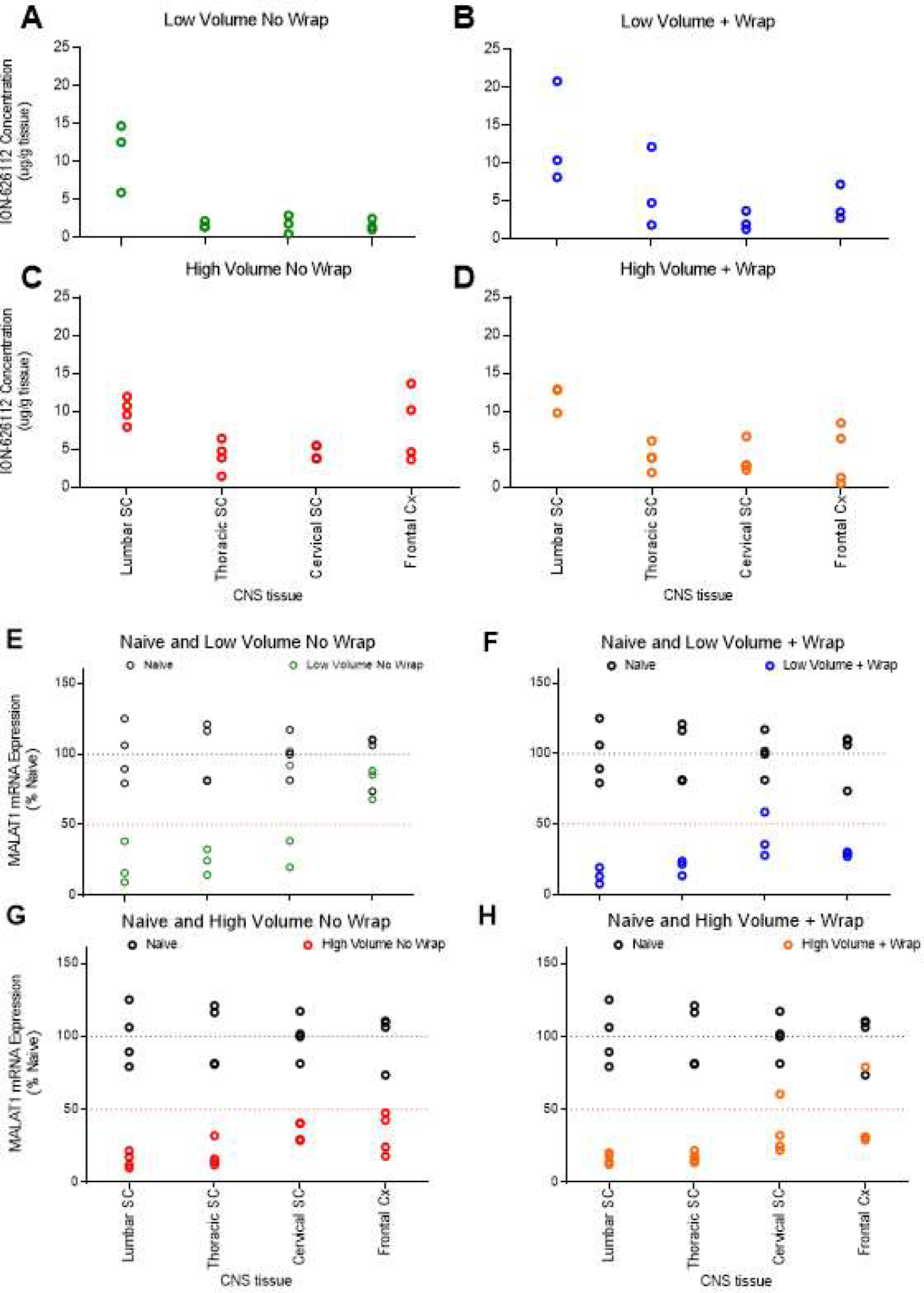
Graphs of the CNS tissue concentrations in µg/g tissue and of the expression levels of *MALAT1* RNA for the animals in the different groups. The CNS tissues are arranged from the tissues closest to the injection site on the left (lumbar spinal cord) to the most distal from the injection site (frontal cortex) on the right. Graph A has data from the animals treated with low volume (0.8 mL) with no percussive wrap treatment in green; graph B has data from the animals treated with low volume with percussive wrap treatment in blue; graph C has data from animals treated with high volume (2.4 mL) with no percussive wrap treatment in red; graph D has data from animals treated with high volume with percussive wrap treatment in orange; graph E has data from the animals treated with low volume (0.8 mL) with no percussive wrap treatment in green; graph F has data from the animals treated with low volume with percussive wrap treatment in blue; graph G has data from animals treated with high volume (2.4 mL) with no percussive wrap treatment in red; graph H has data from animals treated with high volume with percussive wrap treatment in orange. Data in black is the *MALAT1* RNA expression from the naïve animals.

Tissue samples adjacent to those used for the ASO tissue concentration determinations were processed and analyzed for *MALAT1* RNA expression by qRT-PCR. As the MALAT1 ASO is designed to suppression *MALAT1* RNA, this is a measure of pharmacodynamic effect. Consistent with the drug concentrations, *MALAT1* RNA expression is significantly suppressed from naïve expression levels in the spinal cord, but not in the frontal cortex in the low volume and no wrap conditions (Figure 4E, Table 3). However, when the percussive wrap treatment was added to the low volume injection, RNA suppression occurs throughout the CNS structures, including the frontal cortex which is distal from the injection site (Figure 4F, Table 3). With high dose volume alone (Figure 4G) and with high dose volume and percussive wrap treatment (Figure 4H), more uniform suppression of expression of the target RNA can be observed across the CNS with significant reductions of expression compared to naïve animals (Table 3). This pattern of RNA suppression is similar between the two different high dose volume groups (Figures 4G and H, Table 3) suggesting that the addition of percussive forces does not add any distribution benefit to the high dose volume for the distribution of the ASO to rostral structures. The MALAT1 RNA suppression in the frontal cortex in both of the high dose volume treatments are similar to the low dose volume with percussive wrap treatment (Figures 4G, H and F) and dissimilar to the low dose volume without percussive wrap treatment (Figure 4E) suggesting that the addition of percussive wrap increases the rostral distribution of the ASO with low dose volume equal to the increase seen with the high dose volume injection. Taken together, either increasing dose volume, or adding percussive wrap treatment to the low injection volume, could increase rostral distribution of ASOs.

**Table 3.**
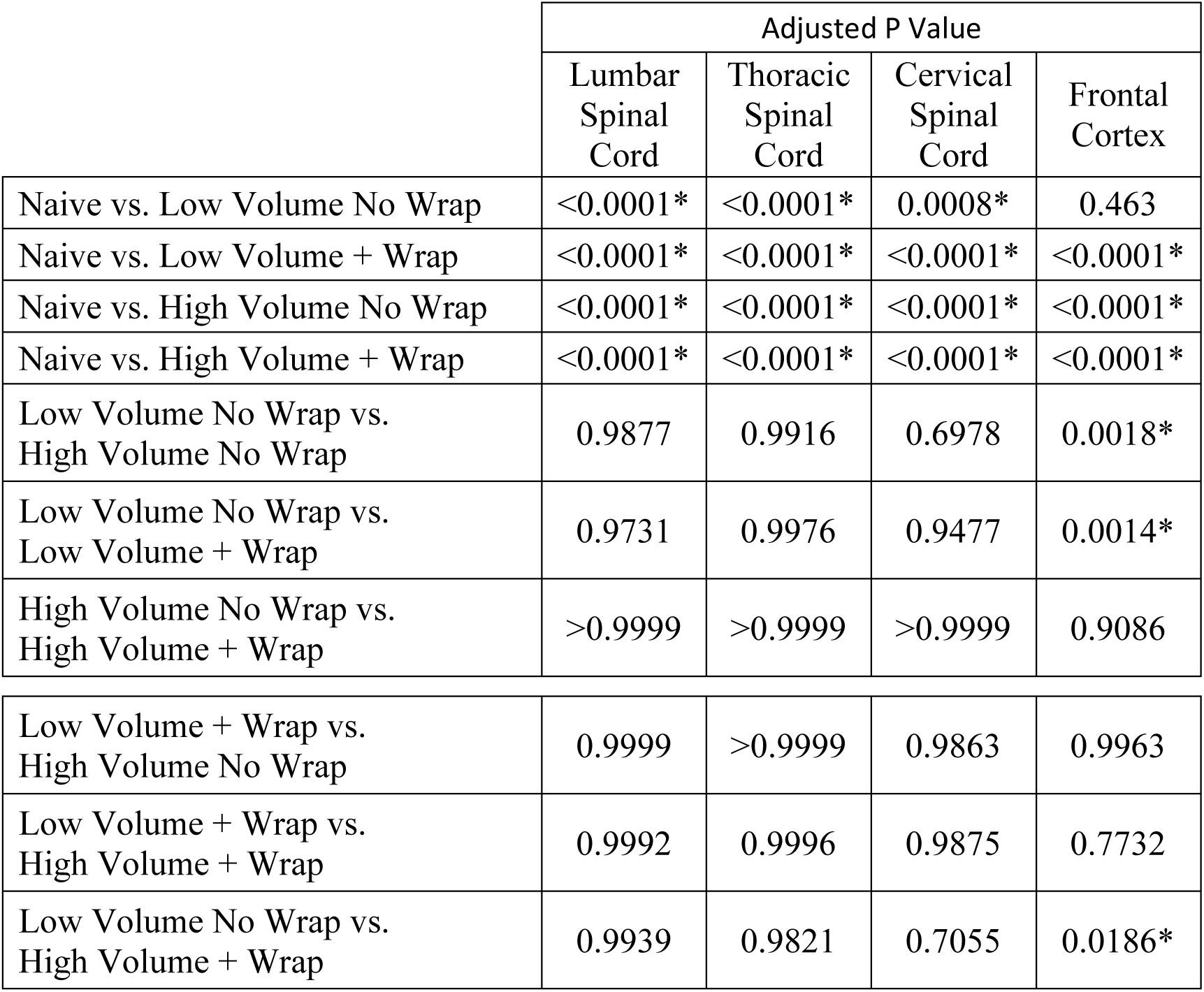
Tukey’s multiple comparisons test for Malat1 RNA expression results

|  | Adjusted P Value |  |  |  |
| --- | --- | --- | --- | --- |
|  | Lumbar Spinal Cord | Thoracic Spinal Cord | Cervical Spinal Cord | Frontal Cortex |
| Naive vs. Low Volume No Wrap | <0.0001* | <0.0001* | 0.0008* | 0.463 |
| Naive vs. Low Volume + Wrap | <0.0001* | <0.0001* | <0.0001* | <0.0001* |
| Naive vs. High Volume No Wrap | <0.0001* | <0.0001* | <0.0001* | <0.0001* |
| Naive vs. High Volume + Wrap | <0.0001* | <0.0001* | <0.0001* | <0.0001* |
| Low Volume No Wrap vs. High Volume No Wrap | 0.9877 | 0.9916 | 0.6978 | 0.0018* |
| Low Volume No Wrap vs. Low Volume + Wrap | 0.9731 | 0.9976 | 0.9477 | 0.0014* |
| High Volume No Wrap vs. High Volume + Wrap | >0.9999 | >0.9999 | >0.9999 | 0.9086 |
| Low Volume + Wrap vs. High Volume No Wrap | 0.9999 | >0.9999 | 0.9863 | 0.9963 |
| Low Volume + Wrap vs. High Volume + Wrap | 0.9992 | 0.9996 | 0.9875 | 0.7732 |
| Low Volume No Wrap vs. High Volume + Wrap | 0.9939 | 0.9821 | 0.7055 | 0.0186* |

To further confirm ASO distribution and pharmacology on a cellular level, spinal cord and frontal cortex histological samples were stained for *MALAT1* RNA via in situ hybridization (ISH) and for ASO via immunohistochemistry (IHC) (Figure 5). The IHC staining for the ASO clearly demonstrates good distribution of the ASO throughout the CNS in the high volume no wrap treated animal and lesser distribution to rostral structures more distal to the injection site in the low volume no wrap treated animal. The *MALAT1* ISH demonstrates complete reduction of RNA expression in all the CNS regions evaluated in the animal from the high volume treated group. In contrast, the *MALAT1* ISH images from the low volume group animal shows complete reduction of RNA expression in the spinal cord regions and only partial reduction of *MALAT1* RNA expression in the frontal cortex. These results are consistent with the ASO concentration, PCR, and imaging data.

**Figure 5:**
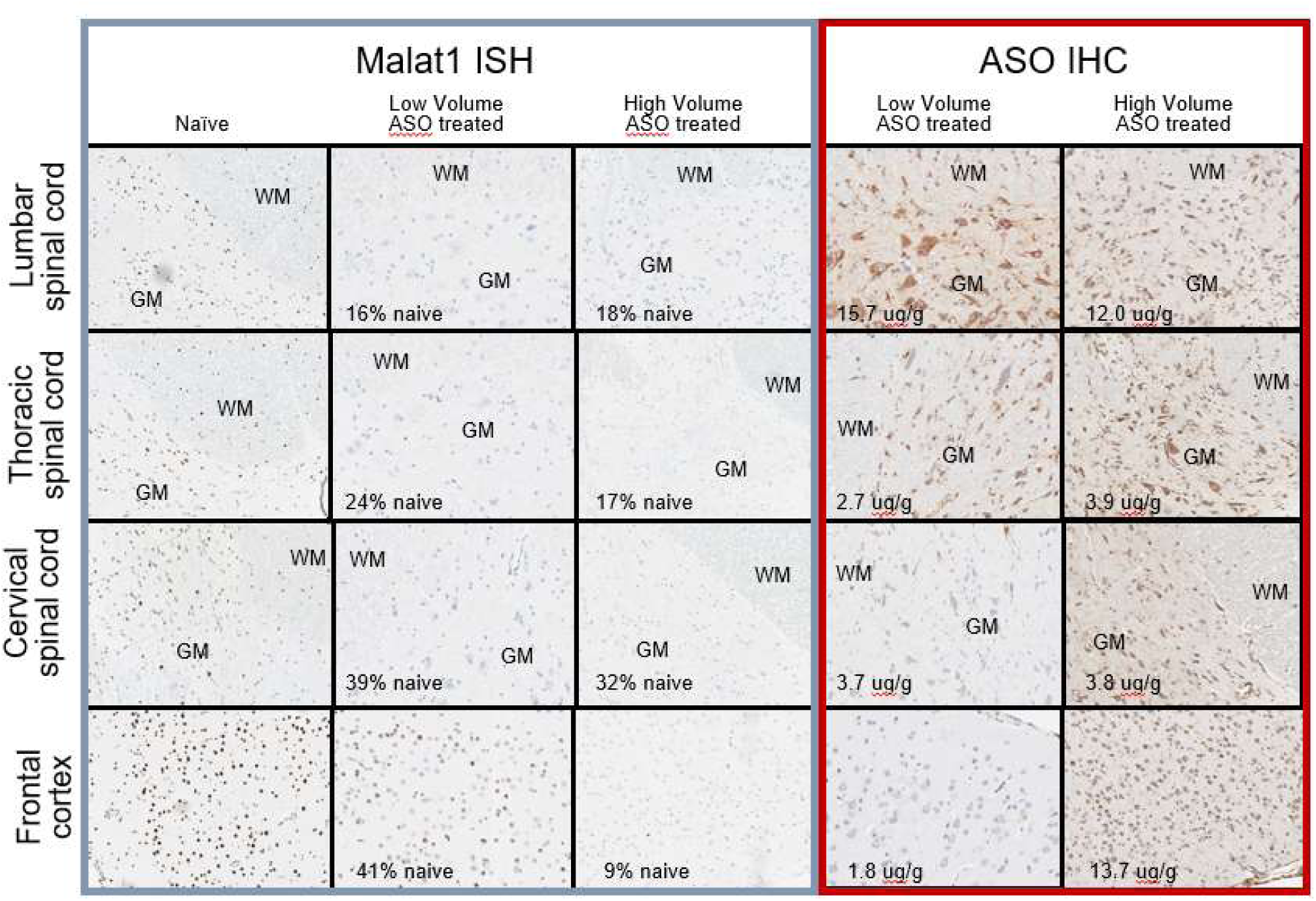
Imaging the spinal cord and frontal cortex ASO distribution and pharmacology. Representative images from *MALAT1* ISH (left three columns) and ASO IHC (two columns on the right) in representative animals from the naïve animal, the low (0.8 mL) volume no wrap, and high (2.4 mL) volume no wrap groups. Images are arranged top to bottom with the proximal to distal areas from the injection site. In the spinal cord pictures, white matter is designated by WM and gray matter by GM. The tissue concentrations in ug/g tissue measured by LC-MS in the adjacent tissues are displayed in the ASO IHC pictures. The *MALAT1* RNA expression values (percent of naïve control) as measured by PCR from the adjacent spinal cord tissues are written on the ASO treated *MALAT1* ISH pictures.

### Broad pharmacology following high volume delivery of a MAPT-ASO

Suppression of human *MAPT* has been proposed as a potential therapy for tauopathies, including Alzheimer’s disease and frontotemporal dementia (8). Indeed, ASOs suppressing *Mapt* in rodent models of disease can reverse pathology and ameliorate phenotype (8). Since many of the potential indications for a MAPT ASO likely require cortical suppression, we evaluated whether we could increase distribution and pharmacology of a MAPT ASO that targets monkey *MAPT* mRNA by using a high volume (2 mL) compared to a low dose volume (0.8 mL). We found a trend toward increased ASO concentrations in the more rostral structures in the high-volume group compared to the low volume group (Figure 6A). Consistent with the ASO concentration data, increasing the dose volume significantly increased the pharmacological action of the ASO, with greater reduction in *MAPT* mRNA in the high-volume group compared to the low volume group (Figure 6B). Taken together, using a therapeutically relevant modality, increasing the convective forces within the CSF following an IT drug delivery can increase ASO concentration and pharmacological action throughout the brain.

**Figure 6.**
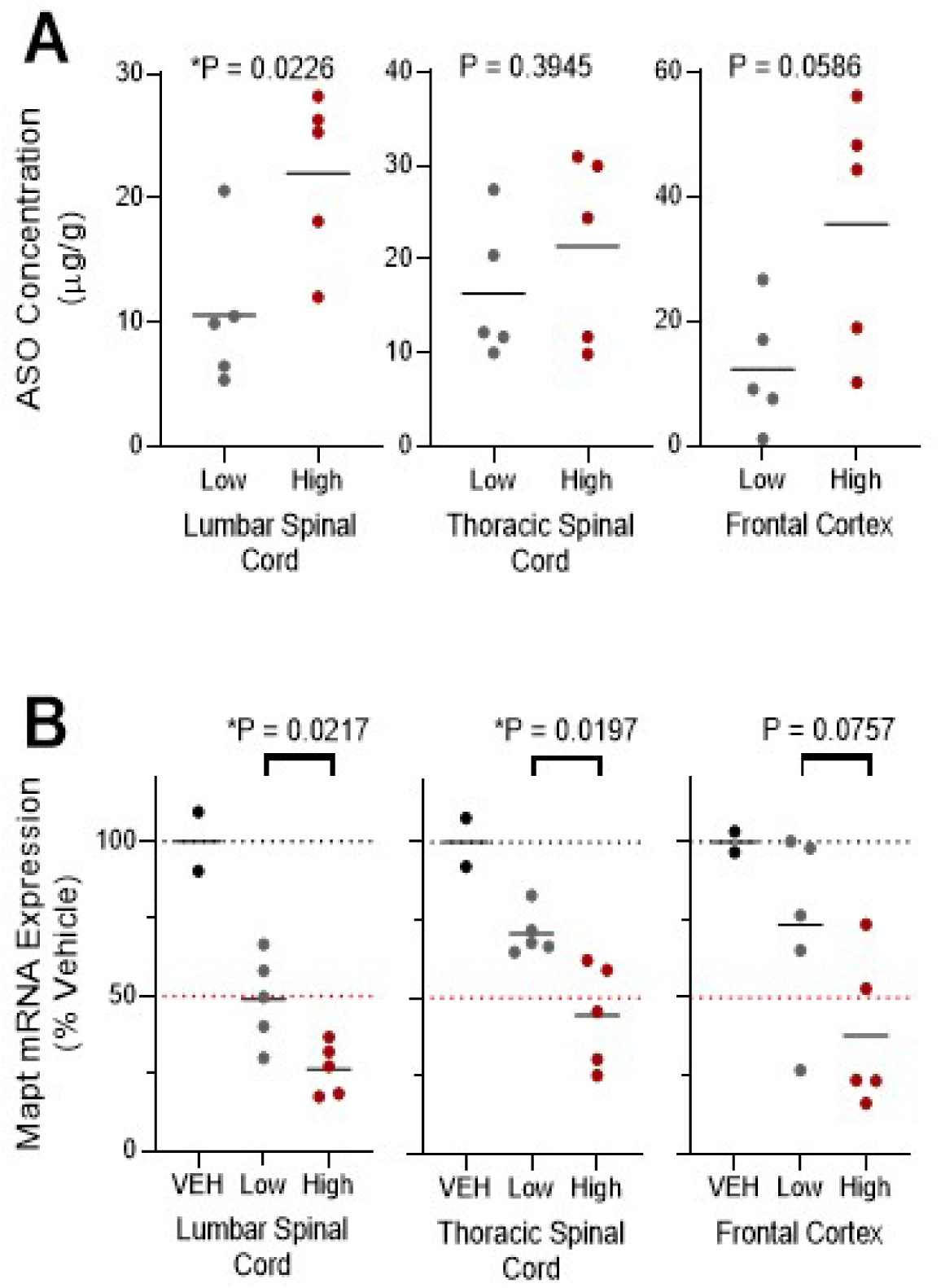
Effect of injection volume on the CNS distribution and pharmacology of an ASO specific to the *MAPT* gene. A) Graphs of the tissue concentrations (µg/g tissue) of the MAPT ASO in the lumbar and thoracic spinal cord, and frontal cortex for the animals treated with low (0.8 mL, gray) or high (2.0 mL, red) dose volume. B) Graphs for the *MAPT* mRNA expression for the low (gray) versus high (red) volume treatments as a percentage of the vehicle treated (black) animals. At the top of each graph is the P value for the comparison of the low versus high volume treatment as calculated with two tailed Welch’s T-test. A star (*) designates a P value less than 0.05. Data is depicted as individual values with horizontal bar depicting the mean value for the group.

## DISCUSSION

We embarked on this series of experiments to evaluate how manipulation of convective forces affected the rostral distribution of drug after lumbar intrathecal administration in the monkey. The objective of this work is to incorporate the lessons learned here in clinical practice to potentially increase the cranial distribution of IT delivered therapeutics. Our earlier work in rodents demonstrated that adjusting the volume of injection can increase neuraxial exposure distal to the lumbar IT injection site (7). There is a small body of literature that describes the effects of several convective mechanisms in the CNS space on CSF motion including CSF turnover, cardiac motion, respiratory thoracic motion, and body movement (3-6). We chose injection volume and exogenous forces applied with a percussive wrap as potential modifiers of ASO distribution, as these would be easily incorporated into the clinical environment.

Although increasing dose volume has not been systematically taken advantage of in IT drug therapy, injected volumes of up to 50 mL (approximately 33% of the total CSF volume) have previously been given safely in humans. Moreover, myelography contrast injection guidelines call for up to 17 mL IT injection of CT contrast media in adults (9, 10).

In our first experiment, ^64^Cu-DOTA was successfully injected IT and imaged in cynomolgus monkeys in high (1.8 mL) or low (0.4 mL) bolus volumes, with or without exogenous force applied to the thorax via a percussive wrap. There was an increase of rostral radioactivity distribution distal to the injection site with the larger bolus dose volume compared to the lower dose volume. Rostral radioactivity distribution was also increased with the addition of the percussive wrap in both the low and high dosing volumes.

These results then led to our next experiment dosing a pharmacological dose of an unlabeled ASO to the non-protein coding RNA *MALAT1* spiked with ^99m^Tc-DTPA to bridge the previous radioactive imaging results with ASO distribution using dose volume and percussive wrap. We again found that the larger injection volume increased rostral distribution of both the radioactivity and the MALAT1 ASO distal from the lumbar injection site. We also found that percussive wrap treatment increased the rostral distribution of the radioactivity and MALAT1 ASO above that seen with the low dose volume. However, percussive wrap did not seem to increase rostral distribution of the ASO or radioactivity above that seen with the high dose volume.

We then utilized the lessons learned from these two experiments and evaluated the distribution of an ASO against *MAPT* after lumbar intrathecal administration dosed in a low (0.8 mL) or high (2 mL) volume. We again found an increase in rostral distribution of the *MAPT* ASO distal from the lumbar injection site with the larger volume when compared to the smaller volume. The rostral distribution of the *MAPT* ASO to the cranium is important because this target is important in frontotemporal dementia and Alzheimer’s disease, both neurodegenerative diseases of the brain. These convective forces effects were also evident in the increase in the action of the ASO in the large dose volume animals compared to the low dose volume ones.

It is likely that both increased dose volume and mechanical percussion involve convection enhancement. In addition to increasing radioactive tracer concentrations in the more cranial regions of the neuraxis, the percussive wrap reduced tracer concentration in the lumbar region in the low volume condition. The use of increased dose volume and exogenous mechanical force may thus be useful for increasing IT therapeutic dose to cranial regions and reducing exposure at the lumbar site of delivery when required. Conversely, for IT dosed drugs targeted for local lumbar spinal cord or nerve root action where neuraxial spread is not desired, it may be useful to minimize convective force by using lower dose volumes.

We evaluated the CSF volume in a large group of cynomolgus monkeys using T2-weighted volumetric MRI methods. We found that the cynomolgus monkey has a total CSF volume of 11.9 ± 1.6 mL (mean ± standard deviation). This value was used to guide selection of IT bolus injection volumes to model published high and low intrathecal injection volumes delivered in the clinic.

Total CSF volume in humans is generally quoted as approximately 130 mL (12). This number is derived from Dixon and Halliburton in 1916 (13) that stated the total CSF volume in humans to be 100-130 mL based on the “estimations of several observers”. Recently, Chazen et al. (14) published a paper using T2-weighted volumetric MRI methods in the human similar to what we employed here. They found a volume of 263 ± 55 mL (mean ± standard deviation) in a group of 15 normal human subjects, which is much larger than the traditional 130 mL volume. When we re-evaluate our injection volume percentages of total CSF volume using these new human data, they change from 3-7% to 1.4-3% for the low volume doses and from 15-20% to 7-10% for the high-volume doses. When allometrically scaling intrathecal dosing between monkey to human, what was previously a 130/11.9 mL, or an approximately 10-fold higher CSF volume in the human, now becomes a 263/11.9 mL or approximately 20-fold higher CSF volume in the human than in the monkey.

Clear assessment of the variability in the quality of IT ^64^Cu-DOTA delivery by injection as revealed by PET imaging also illustrates the value of molecular imaging in assessing efficiency of IT therapeutics delivery. IT nuclear imaging, which has long been used to evaluate CSF leaks, may provide great value in understanding the efficiency and pharmacodynamics of IT dosed therapeutics. Experiments in humans are required to assess the translatability of the volume principles demonstrated by the experiments within this manuscript. The assumption that rostral distribution as a function of percentage of CSF volume injected is consistent from NHP to human has yet to be tested. Nuclear imaging will allow us to test these volume principles in humans. It may also be crucial in determining differences in intrathecal dynamics in different disease states, for instance where cerebral ventricular volumes are increased, or where cortical atrophy increased subarachnoid CSF space.

## ABBREVIATIONS

IT: Intrathecal
CNS: Central Nervous System
CSF: Cerebral spinal fluid
ASO: Antisense oligonucleotide
NHP: nonhuman primate
MRI: Magnetic Resonance Imaging
PET: Positron Emission Tomography
SPECT: Single Photon Emission Computed Tomography
CT: Computed Tomography
cm^3^: cubic centimeters
BBB: Blood Brain Barrier
DOTA: dodecane tetraacetic acid
DTPA: diethylenetriaminepentaacetic acid
MAPT: microtubule associated protein tau
NBR: Northern Biomedical
Malat1: metastasis associated lung adenocarcinoma
SC: subcutaneous
IM: intramuscular
FSE: fast spin echo
ROI: region of interest
HFCWO: high-frequency chest wall oscillation
ITLC: instant thin layer chromatography
OSEM2D: Ordered Subset Expectation Maximization
%ID/g: percent injected dose per gram
MOE: 2’ methoxyethyl
aCSF: artificial cerebral spinal fluid
MLEM: maximum likelihood estimation method
QF: quantification factor
MIP: Maximum intensity projection
qRT-PCR: quantitative reverse transcriptase polymerase chain reaction
LC-MS/MS: liquid chromatograph with mass spectrometry detection
TEA: triethyalamine
HFIP: hexo-fluoro-isopropanol
IS: internal standard
ACD: Advanced Cell Diagnostics
DAB: 3,3’-diaminobenzidine chromogen
AUC: area under the curve
ISH: in situ hybridization
IHC: immunohistochemistry
CV%: Percent coefficient of variation
SD: Standard Deviation

## DECLARATIONS

### Ethics Approvals

The MRI quantification of CSF volume experiment in the monkey, ^64^Cu-DOTA and ^99m^Tc-DTPA/Malat1 ASO experiments were performed at MPI Research and the study procedures were reviewed and approved by the Institutional Animal Care and Use Committee at MPI Research prior to the start of the study. The ^64^Cu-DOTA study was performed under approved protocols 1881-091 and 1881-100. The ^99m^Tc-DTPA/Malat1 ASO experiment was performed under approved protocol 1881-102 and the MRI experiment was performed under protocols 1881-098 and 1881-100. The Mapt ASO experiment was performed at Northern Biomedical Research under a protocol approved by their Institutional Animal Care and Use Committee, specifically protocol number 004-031. All three of these studies complied with the U.S. Department of Agriculture’s Animal Welfare Act (9 CFR Parts 1, 2, and 3) and the Guide for the Care and Use of Laboratory Animals, Institute of Laboratory Animal Resources, National Academy Press, Washington, D.C., 2011. Both MPI Research and Northern Biomedical Research are accredited by the Association for Assessment and Accreditation of Laboratory Animal Care (AAALAC).

### Consent for Publication

Not applicable.

### Availability of Data and Materials

The datasets used and/or analyzed during the current study are available from the corresponding author on reasonable request.

## Authors Contributions

J.M.S and C.M. assisted in designing the experiments and analyzed data and are co-primary authors of the manuscript. C.M. also conducted experiments and collected data. B.P., L.T., B.D-S., and B.F. conducted experiments, collected, analyzed data and assisted in writing the manuscript. R.P., and J.H. analyzed data and assisted in writing the manuscript. D.A.W and L.H. assisted in designing experiments and writing the manuscript. S.H. assisted in designing and conducting the experiments and writing the manuscript. N.C. and H.K. assisted in writing the manuscript and interpreting the data. A.V., and J.H. assisted in designing experiments, analyzed data and assisted in writing the manuscript. E.E.S. assisted in designing the experiments and writing the manuscript and is the senior member of the Ionis group.

## Acknowledgements

The authors would like to acknowledge the scientists, staff, and technicians at MPI Research and Invicro for their precise execution of the studies described in this paper. The authors would like to specifically thank Kevin Magalhaes, Joshua Higgins, and Carolynn Gaut for their hard work and dedication to the project management of the imaging studies and Howard Dobson for useful discussions and his valuable insight. The authors would also like to acknowledge Tracy Reigle and Wanda Sullivan for their assistance with the figures in this publication.

